# Application of seaweed organic components increases tolerance to Fe deficiency in tomato plants

**DOI:** 10.1101/553917

**Authors:** Sandra Carrasco-Gil, Raúl Allende-Montalbán, Lourdes Hernández-Apaolaza, Juan José Lucena

**Author notes:** (S. Carrasco-Gil); (JJ Lucena).

## Abstract

The beneficial effects of seaweed extracts have been related to plant growth regulators present in seaweeds. However, algae extracts comprise other organic compounds such as phenols, mannitol, alginates, laminarins and fucoidans that may have a relevant role regarding abiotic stress tolerance due to Fe deficiency. Therefore, we evaluated the individual effect of these organic compounds on the mitigation of Fe deficiency applying a range of concentrations (x1/10, x1, x10) in agar Petri dishes (in tomato seeds) and in the nutrient solution of a hydroponic system (tomato plants). Germination and plant growth promotion, root morphology, chlorophyll content and antioxidant activity were determined. Results showed that the lowest concentration x1/10 and phenolics, laminarin and fucose compounds contributed to increase the tolerance to Fe deficiency in tomato plants.

## Introduction

The use of commercial seaweed extracts (SWEs) in agriculture is an increasingly widespread practice, since these products may enhance the plant growth development and the tolerance to abiotic stresses, which are increasing due to climate change [1–3]. However, the information on the mode of action of these extracts is scarce. This fact, along with the high variability in the composition of these products, may reduce the confidence that farmers have on the SWEs formulations. Different research works that studied SWEs obtained from the same type of seaweed source (as example *A. nodosum*), showed different results [4] probably due to the variability in the composition among batches, extraction process used for their manufacturing, and the doses, frequency and time of application to crops.

The forthcoming Europe legislation regarding fertilizer products (*REGULATION OF THE EUROPEAN PARLIAMENT AND OF THE COUNCIL laying down rules on the making available on the market of CE marked fertilising products and amending Regulations (EC) No 1069/2009 and (EC) No 1107/2009*) does not establish requirements regarding algae extract components, except the maximum allowed concentration of several contaminants that may be present in algae extract products. However, algae extracts comprise many organic compounds such as betaines, proteins, phenols, vitamins, vitamin precursors, plant growth regulators, mannitol, alginates, laminarins and fucoidans [5], which are not taken into account in the proposed EC regulation, and that may have a relevant role regarding growth promotion and abiotic stress tolerance.

The European Food Safety Authority [6] has related alginates, fucoidan and mannitol to the effects of SWE, mainly because these compounds make up the majority of the organic composition of these extracts. Alginates can improve soil conditions promoting the formation of aggregates between soil particles and, therefore, increasing the absorption and translocation of nutrients [7,8], root growth and soil microbial activity [9]. Mannitol has several functions within plant systems, being able to act as a reserve carbohydrate [10] or as a protective agent against reactive oxygen species (ROS) [11]. Some authors have observed an increase in enzymatic antioxidant activity after the application of mannitol in plants under salt stress [12]. In addition, the application of mannitol can improve the protection of the roots against lipid peroxidation [13]. Fucoidans may promote the antioxidant activity [14,15], although it has not yet been tested on plants. However, there are also other organic compounds such as phenolics or laminarin, relatively abundant in the SWEs whose presence has been related to some of its effects. The phenolic compounds are particularly abundant in brown seaweed and are known for their antioxidant activity. Among the phenolic compounds, salicylic acid (SA) can alleviate the effects caused by various abiotic stress factors, such as extreme heat [16], soil salinity [17,18] or drought [19]. Gallic acid (GA) is another important phenolic compound, which improved the plant tolerance to abiotic stress such as soil salinity [20], ozone exposure [21] and low temperature [22]. Laminarin can account for 0-33% of the total dry weight of the marine algae used in SWE production [23]. Laminarin can modulate the antioxidant system of chloroplasts in situations of abiotic stress [24].

There is scarce information about the effect of SWEs on nutrient deficiency. Several studies revealed that SWE application might stimulate nutrient uptake and translocation in plants [25–28]. Iron (Fe) is an essential nutrient in plant nutrition that is involved in chlorophyll (Chl) synthesis, in electron transport photosynthesis, DNA or hormone synthesis and N-fixation process [29]. Iron can also act as a cofactor within many antioxidant enzymes, such as catalase (CAT) and superoxide dismutase (SOD), which are responsible for protection against reactive oxygen species (ROS) production [30,31]. A low Fe availability, especially in calcareous soils with alkaline pH, results in a reduction of plant productivity and quality [32]. Nutrient imbalances such as Fe deficiency may be alleviated by the use of SWE products, enhancing the defense mechanism to reduce the oxidative stress and the chlorosis. Moreover, the addition of SWEs may promote the root development and the photosynthesis, improving the nutrient uptake.

Therefore, the objective of this work was to study the effect of individual application of organic components present in the algae extracts on: (1) plant growth development and (2) mitigation of abiotic stress due to Fe deficiency (nutrient imbalance). Also, (3) identify the concentration range at which these compounds may have positive effects. This work will contribute to scientific basis for establishing criteria for the production, use, and regulation of new seaweed extract products, which guarantee farmers the benefits indicated on the label package of these type of products.

## Material and Methods

### Organic compounds

A set of organic compounds present in SWEs that have been related to the beneficial effects of the SWE application in agriculture by several authors (see introduction section) were selected: Fucose (L-(-)-Fucose, Sigma-Aldrich) that is the fundamental sub-unit of the fucoidan; alginic acid polysaccharide (alginic acid sodium salt, Sigma-Aldrich), laminarin polysaccharide (from *Laminaria digitate*, Sigma-Aldrich); mannitol that is a sugar alcohol (D-Mannitol, Merck); salicylic acid (Panreac) and gallic acid (Sigma-Aldrich) that are phenolic compounds. The organic compounds were applied at three concentrations x1/10, x1, x10 (Table 1). The concentration x1/10 (1/10-fold with respect to x1) and x10 (10-fold with respect to x1). The concentration *x*1 was calculated based on the concentration of these organic compounds in commercial SWEs of *A. nodosum* [4,33] and the average dose of application (by root) of commercial SWEs for tomato plants. Control treatment without any organic compound application was also performed.

**Table 1.** Organic compound concentration (mg/L) of salicylic acid (SA), gallic acid (GA), sodium alginate (SoA), mannitol (MA), laminarin (LA) and fucose (FU) applied in seeds and tomato plants.

| Organic compound | Concentration (mg/L) |  |  |
| --- | --- | --- | --- |
|  | x1/10 | x1 | x10 |
| SA | 4,7 | 47 | 470 |
| GA | 4,7 | 47 | 470 |
| SoA | 9,0 | 90 | 900 |
| MA | 5,8 | 58 | 580 |
| LA | 0,070 | 0,70 | 7,0 |
| FU | 3,1 | 31 | 310 |

### Germination assay in Petri dish

Firstly, sterilized Petri dishes were prepared for seed germination adding individually the organic compounds (SA, GA, SoA, MA, LA, FU) at three concentrations (x1/10, x1 and x10; see Table 1) in 1.5% (w/v) plant agar (Duchefa Biochemia, Haarlem, Netherlands). Tomato seeds (*Solanum lycopersicum* L. Moneymaker) were surface sterilized, vernalized at 4 °C for 24 h in darkness and placed on Petri dishes (8 seeds per Petri dish) with the respective organic compounds treatments and concentrations. Control treatment without any organic compound application was also performed. The sowing was carried out in a laminar air flow cabinet to avoid bacterial contamination. Petri dishes were placed in vertical position with a slight inclination for 7 days in a growth chamber with a photosynthetic photon flux density at leaf height of 1000 μmol m^-2^ s^-1^ photosynthetically active radiation, 16-h, 25 °C, 40% humidity/ 8-h, 20 °C, 60% humidity day/night regime. A total of 2 Petri dishes per treatment and concentration was performed. The germination percentage (%) was measured at day 2 and the growth promotion (+%) or growth inhibition (-%) root seedlings at day 3, 4, 5 and 7 after sowing. The growth promotion and growth inhibition were calculated as follow:

Growth (%) = [(treated root length – control root length)/control root length]×100

### Plant material and growth conditions

Tomato *(Solanum lycopersicum cv.* Moneymaker) plants were grown in a growth chamber with a photosynthetic photon flux density at leaf height of 1000 μmol m^-2^ s^-1^ photosynthetically active radiation, 16-h, 25 °C, 40% humidity/ 8-h, 20 °C, 60% humidity day/night regime. Seeds were surface sterilized and germinated in vermiculite for 16 days in 1/20 diluted Hoagland nutrient solution in distilled water. Seedlings were pre-adapted to hydroponic system in 10-L boxes (28 plants per box) in 1/5 diluted Hoagland nutrient solution with 20 μM Fe and pH 6.0 during nine days. Plants were them transferred to 100 mL plastic pots (one plant per pot) and grown in completed Hoagland solution containing in mM: 7.5 NO_3_^−^, 1.0 H_2_PO_4_^−^, 1.05 SO_4_^-2^, 3.5 K^+^, 2.5 Ca^2+^, 1 Mg^2+^; and in μM: 23.2 B, 4.6 Mn^2+^, 1.2 Zn^2+^, 0.18 Cu^2+^, 4.6 Cl^−^, 0.12 Na^+^, 0.12 MoO_4_^2-^ with 20 μM Fe-HBED (Fe(III)-ácido N,N’-bis(2-hidroxibencil)etilendiamino-N,N’diacético) during 10 days. The pH was fixed at around 7.5 by the addition of 0.1 mM HEPES and 0.1 g/L CaCO_3_ to simulate calcareous conditions. The nutrient solution was renewed every 5 days. After that, plants were transferred to 300 mL plastic pots and Fe deficiency was induced removing the Fe-HBED from the nutrient solution with a pH of 7.5 during 6 days.

Organic compounds treatments (salicylic acid (SA), gallic acid (GA), sodium alginate (SoA), mannitol (MA), laminarin (LA) and fucose (FU)) were applied three times during the experiment at two different concentrations (*x*1/10 and *x*1). The concentration *x*10 produced inhibition of germination and growth development in seedling during the Petri dish assay, so that concentration was dismissed in hydroponic experiment. First application of the organic compounds was the first day of growth period with completed Hoagland solution, the second application was after five days of growth period with the renewal of nutrient solution, and the third application was at the beginning of Fe-deficient period. A control treatment without organic compound application was also performed. A total of 4 pots per treatment and concentration were used.

### Plant analysis

The morphology of tomato roots treated with the organic compounds was analysed in Fe sufficiency and after 6 days of Fe deficiency. Fresh roots were washed and blotted with filter paper. Then, root tips and root pieces (3 cm length) cut at five cm from the root tip were mounted on microscope glass slides and analysed by a stereomicroscope (Leica MZ12.5, Wetzlar, Germany) connected to a video camera (Olympus UC30, Tokio, Japan). Leaf chlorophyll index was assessed at day 0, 3 and 5 of Fe deficiency using a SPAD 502 apparatus (Minolta Co., Osaka, Japan) after applying the organic compounds at two concentrations. Data was the average of five measurements of new leaf levels during the Fe-deficient period in a total of four plants per treatment. At the end of the experiment, after 6 days of Fe deficiency, plants (41 day-old) were collected and washed with distilled water. Then, the plan material was divided in root, stem, developed leaves and new leaves and fresh weight (FW) was determined. After that, plant material was stored at −80°C for oxidative stress analysis. Enzymes were extracted from 0.1 g of intact frozen roots and leaves with 1 mL extraction solution, freshly prepared containing 50 mM potassium phosphate buffer at pH 7.8, 2 mM Na_2_-EDTA (ethylene diamine tetraacetic acid), 10 mM DTT (1,4-Dithiothreitol), 20 mM ascorbic acid, 0.6% PVPP (polyvinyl polypyrrolidone) and 50 µl protease inhibitors cocktail. The extracts were centrifuged at 14,000 × g for 15 min at 4°C, and the supernatants were used for the enzymatic assays. Total superoxide dismutase activity (SOD; EC 1.15.1.1) was assayed according to Giannopolitis and Ries [34] with some modifications. Briefly, 300 µl reaction mixture containing 50 mM potasium phosphate buffer pH 7.8, 0.1 mM Na_2_-EDTA, 13 mM methionine, 2 mM riboflavine and 75 mM NBT (nitroblue tetrazolium) were added to 10 µl of crude extract in a microplate. The reaction started by exposing the mixture to cool white fluorescent light and absorbance at 560 nm was measured at 0, 15 and 30 min using a spectrophotometer (Spectro start nano, BMG Labtech, Germany). One unit of SOD activity was defined as the amount of enzyme that causes 50% NBT reduction by superoxide radicals, and the specific activity was expressed as units mg^-1^ of protein. Catalase activity (CAT, EC 1.11.1.6) was determined according to Aebi [35] with some modifications. CAT activity was assayed in a 3 mL reaction volume at 25 °C by adding 0.1 mL of diluted extract to a solution containing 50 mM phospate buffer pH 7.0 and 10 mM H_2_O_2_ The activity was measured by monitoring the decrease in absorbance at 240 nm as a consequence of H_2_O_2_ consumption using a spectrophotometer (Spectro start nano, BMG Labtech, Germany). Activity was expressed as units (mmol of H_2_O_2_ decomposed per minute) per mg of protein.

### Statistical analyses

Statistical analysis was carried out with SPSS for Windows (v. 21.0), using a Levene test for checking homogeneity of variances, and ANOVA or Welch’s tests (*p* < 0.10) were performed. *Post hoc* multiple comparisons of means were carried out using Duncan’s or Games-Howell’s test (*p* < 0.10) as appropriate.

## Results

### Germination assay in Petri dish

In general, the individual application of the organic compounds at different concentration (*x*1/10, *x*1 and *x*10) in tomato seeds showed a higher germination (>6-37 %) with respect to untreated control, except in the case of SA (*x*1 and *x*10) and SoA (*x*1) that was similar to untreated control (Table 2). The GA (*x*1/10) treatment showed the highest germination (37%) compared to the control, followed by GA (*x*1), SoA (*x*10), MA (*x*1 and *x*10) with a 25% of germination. After the seed sowing and organic compounds application, length of root seedlings was measured at day 3, 4, 5 and 7, and compared to the control. The growth promotion and growth inhibition were calculated. The application of organic compounds promoted the root growth during the 7 days compared to the control, with the exception of SA (x10), GA (x10) and FU (x1/10) that inhibited the root growth (Fig 1). At day 3, GA (x1/10; x1) and LA (x1/10) significantly increased the root length (>40%), but SA (x10) treatment totally inhibited the root growth compared to the control. At day 4 and 5, SA (x10) maintained almost total inhibition of root growth compared to the control, and at day 7, SA (x10) and also GA (x10) inhibited the root growth (80% and 40% respectively) with respect to the control.

**Table 2.** Germination (%) at day 2 after organic compounds (salicylic acid (SA), gallic acid (GA), sodium alginate (SoA), mannitol (MA), laminarin (LA) and fucose (FU)) application at three concentrations (x1/10; x1 and x10). A control (C) treatment without organic compound application was performed.

| Organic compound | Germination (%) |  |  |  |
| --- | --- | --- | --- | --- |
|  | C | x1/10 | x1 | x10 |
| **SA** | 0 | 6 | 0 | 0 |
| **GA** | 0 | 37 | 25 | 6 |
| **SoA** | 0 | 12 | 0 | 25 |
| **MA** | 0 | 12 | 25 | 25 |
| **LA** | 0 | 12 | 6 | 19 |
| **FU** | 0 | 6 | 6 | 6 |

**Fig 1.**
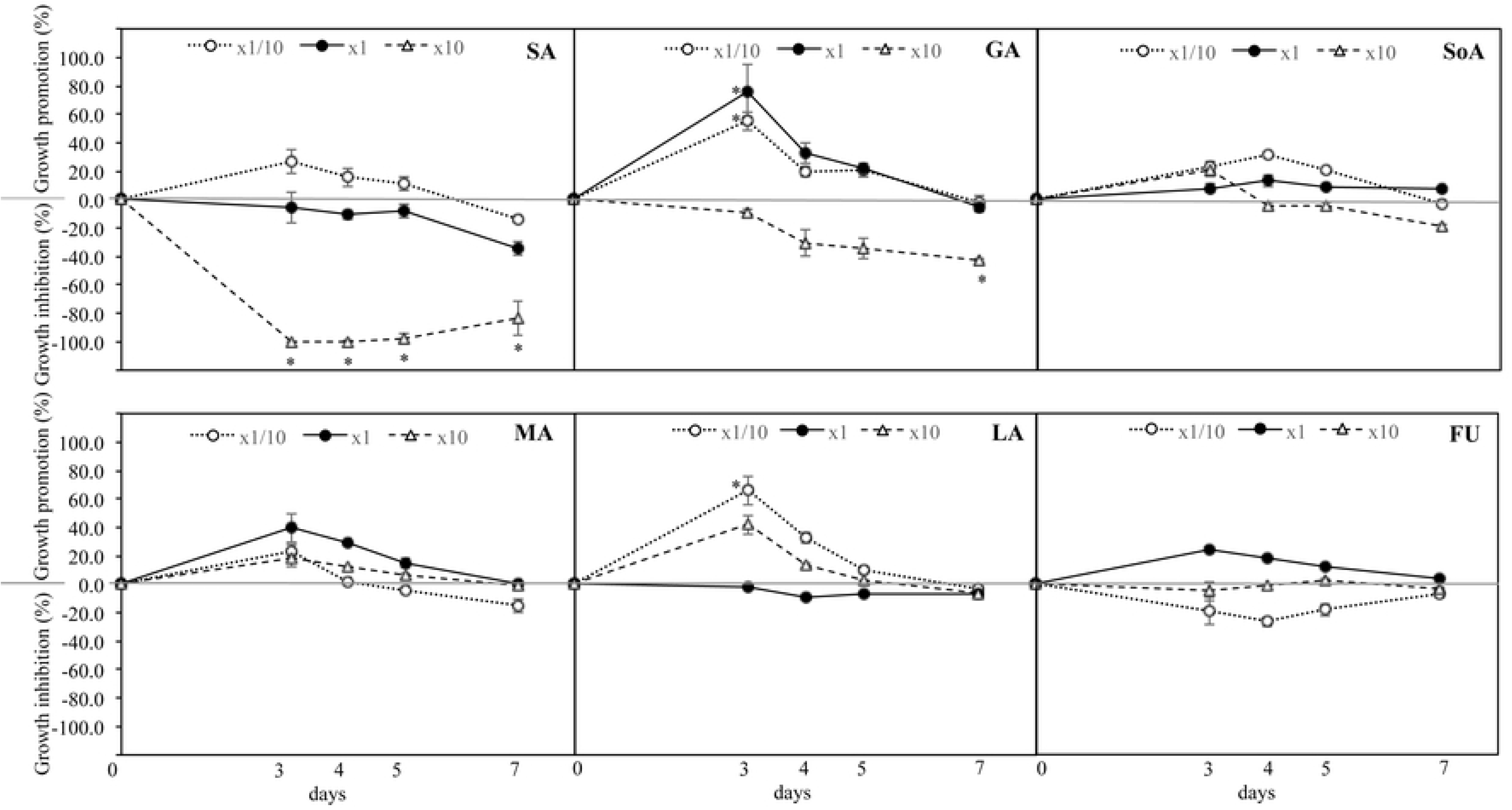
Growth promotion (+%) or growth inhibition (-%) root seedlings at day 3, 4, 5 and 7 after organic compounds (salicylic acid (SA), gallic acid (GA), sodium alginate (SoA), mannitol (MA), laminarin (LA) and fucose (FU)) application at three concentrations (x1/10; x1 and x10) with respect to the untreated control. Data are means ± SE (n=3). Increases or decreases > 40% are indicated by asterisk (*).

### Fresh weight

The FW of root, stem, developed leaves and new leaves was determined after six days of Fe deficiency. The root FW was significantly increased with SA, GA, MA, LA, FU (x1/10) and SoA, MA (x1) compared to the untreated control (Fig 2). The stem FW was significantly increased with SA, GA, LA and FU (x1/10) compared to the untreated control. The developed leaves FW was significantly increased with GA, LA and FU (x1/10) compared to the untreated control. And new leaves FW was significantly increased with all organic compounds (x1/10) and SA, SoA and MA (x1) compared to the untreated control.

**Fig 2.**
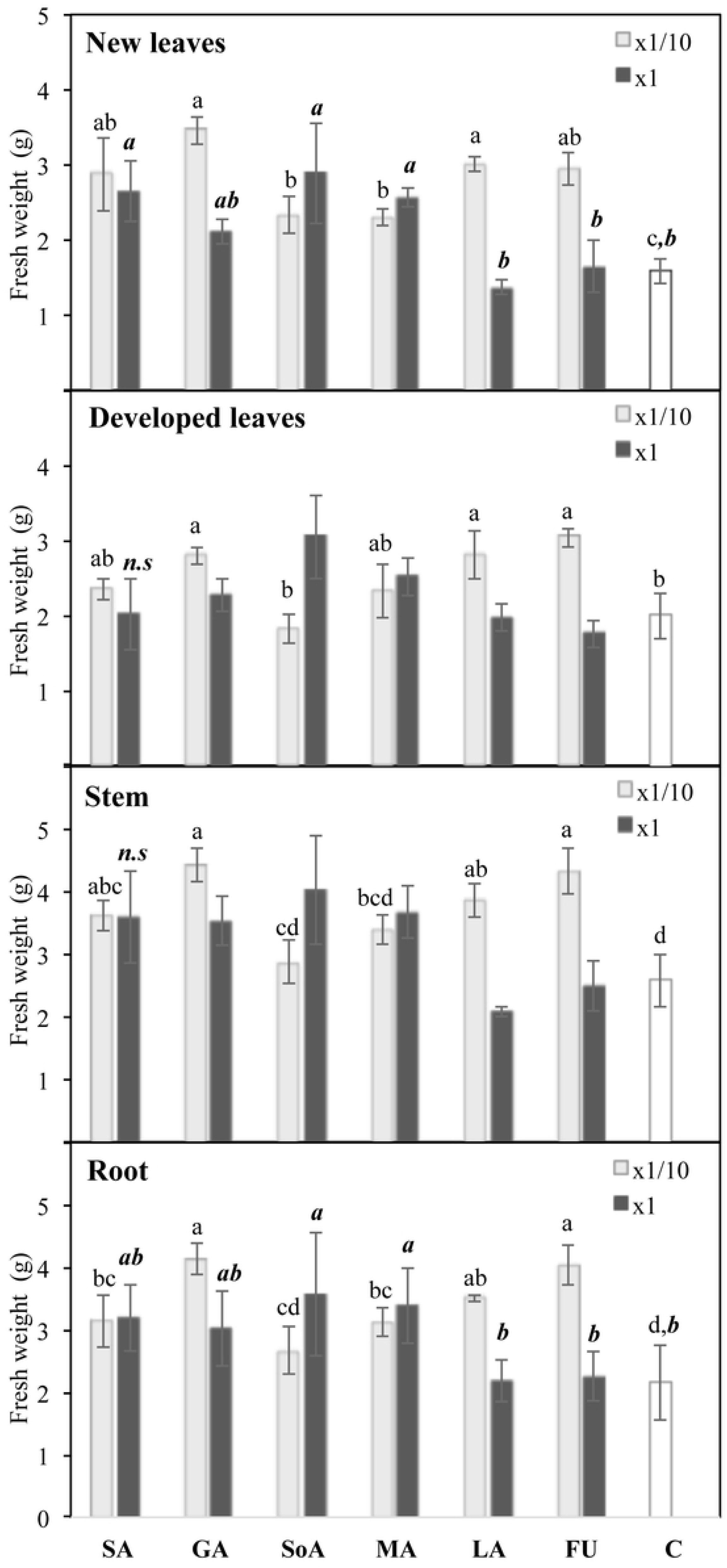
Fresh weight of new and developed leaves, stem and root of tomato plants treated with organic compounds (salicylic acid (SA), gallic acid (GA), sodium alginate (SoA), mannitol (MA), laminarin (LA) and fucose (FU)) at two concentrations (x1/10 and x1) and after 6 days of Fe deficiency. A control (C) treatment without organic compound application was performed. Data are means ± SE (n=3). Significant differences between treatments (*P* < 0.10) are indicated by different letter (regular letters for x1/10; cursive bold letters for x1). Not significant differences between treatments are indicated by n.s.

### Morphology of tomato roots

The morphology of roots treated with organic compounds was evaluated in Fe sufficiency and after 6 days of Fe deficiency compared to untreated control (Fig 3). Under Fe sufficiency, GA, MA (x1) and GA (x1/10) treatments increased the development of root hairs. However SA, GA, MA, FU (x1) and SoA, MA, LA (x1/10) treatments decreased the length of secondary roots with respect to the untreated control. After 6 days of Fe deficiency, LA (x1/10) treatment increased the length of secondary roots, but SoA (x1/10) and GA, LA, FU (x1) decreased the length of secondary roots with respect to the untreated control. Also SA, GA, SoA, MA, FU (x1) and GA, LA, FU (x1/10) treatments increased the development of root hairs with respect to the control. SoA, MA, FU (x1) and SA, SoA, MA, LA (x1/10) treatments increased the distance between secondary roots compared to the control.

**Fig 3.**
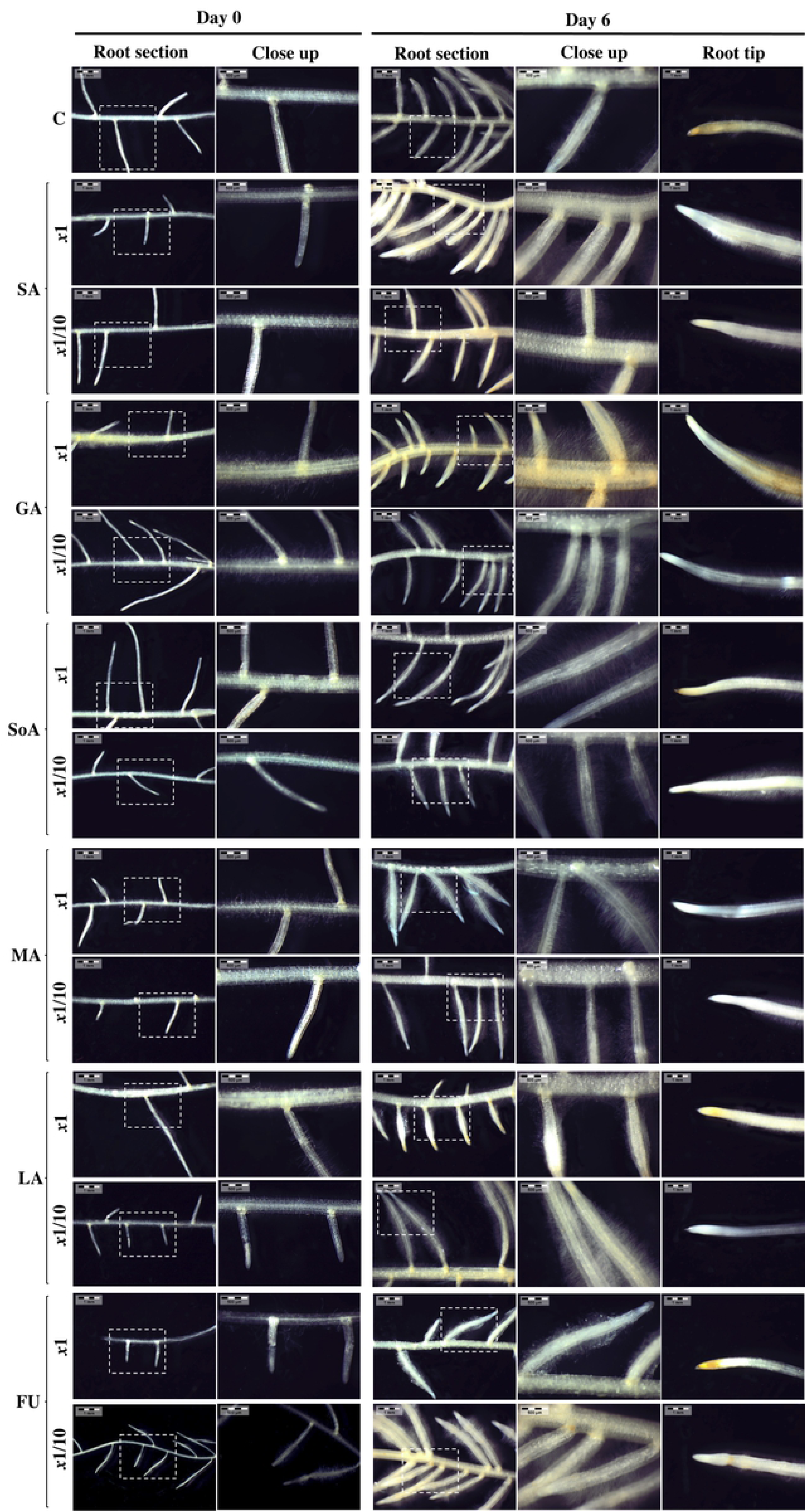
Root morphology of tomato plants treated with organic compounds (salicylic acid (SA), gallic acid (GA), sodium alginate (SoA), mannitol (MA), laminarin (LA) and fucose (FU)) at two concentrations (x1/10 and x1) after 0 and 6 days of Fe deficiency. A control (C) treatment without organic compound application was performed.

### Leaf chlorophyll index

During Fe sufficiency, LA (x1/10) and SA (x1) significantly decreased the chlorophyll index compared to the untreated control, but the rest of treatments maintained the chlorophyll index similar to the control (Fig 4). During Fe deficiency, GA (x1/10) and LA (x1) significantly decreased the chlorophyll index compared to the untreated control at day 3, but FU (x1/10; x1) significantly increased the chlorophyll index compared to the untreated control at day 5 of Fe deficiency.

**Fig 4.**
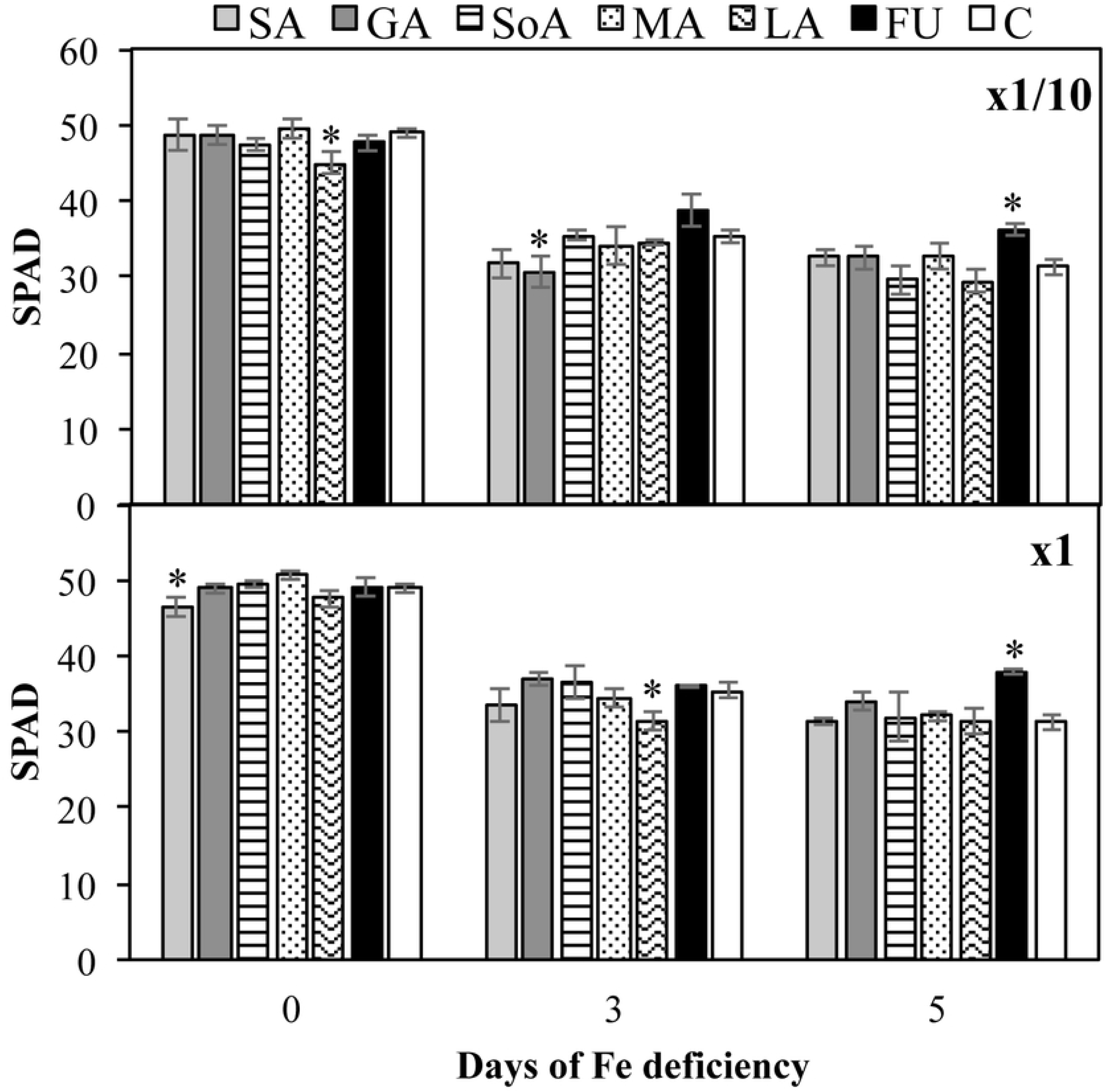
SPAD (chlorophyll index) in new leaves of tomato plants treated with organic compounds (salicylic acid (SA), gallic acid (GA), sodium alginate (SoA), mannitol (MA), laminarin (LA) and fucose (FU)) at two concentrations (x1/10 and x1) at day 0, 3 and 5 of Fe deficiency. A control (C) treatment without organic compound application was performed. Data are means ± SE (n=3). Significant differences between treatments (*P* < 0.10) with respect to the control are indicated by asterisk (*).

### Oxidative stress parameters

The oxidative stress was measured in roots and new developed leaves after six days of Fe deficiency by the determination of SOD and CAT activity. Neither of organic compounds treatments increased the SOD activity in root and new leaves, but SA, GA, SoA and LA (x1) significantly decreased the SOD activity in new leaves compared to the untreated control (Fig 5). Moreover, GA and FU (x1/10) in root, and GA (x1/10) in new developed leaves increased the CAT activity with respect to the control. However, neither of organic compounds treatments significantly decreased the CAT activity in root and new leaves compared to the untreated control.

**Fig 5.**
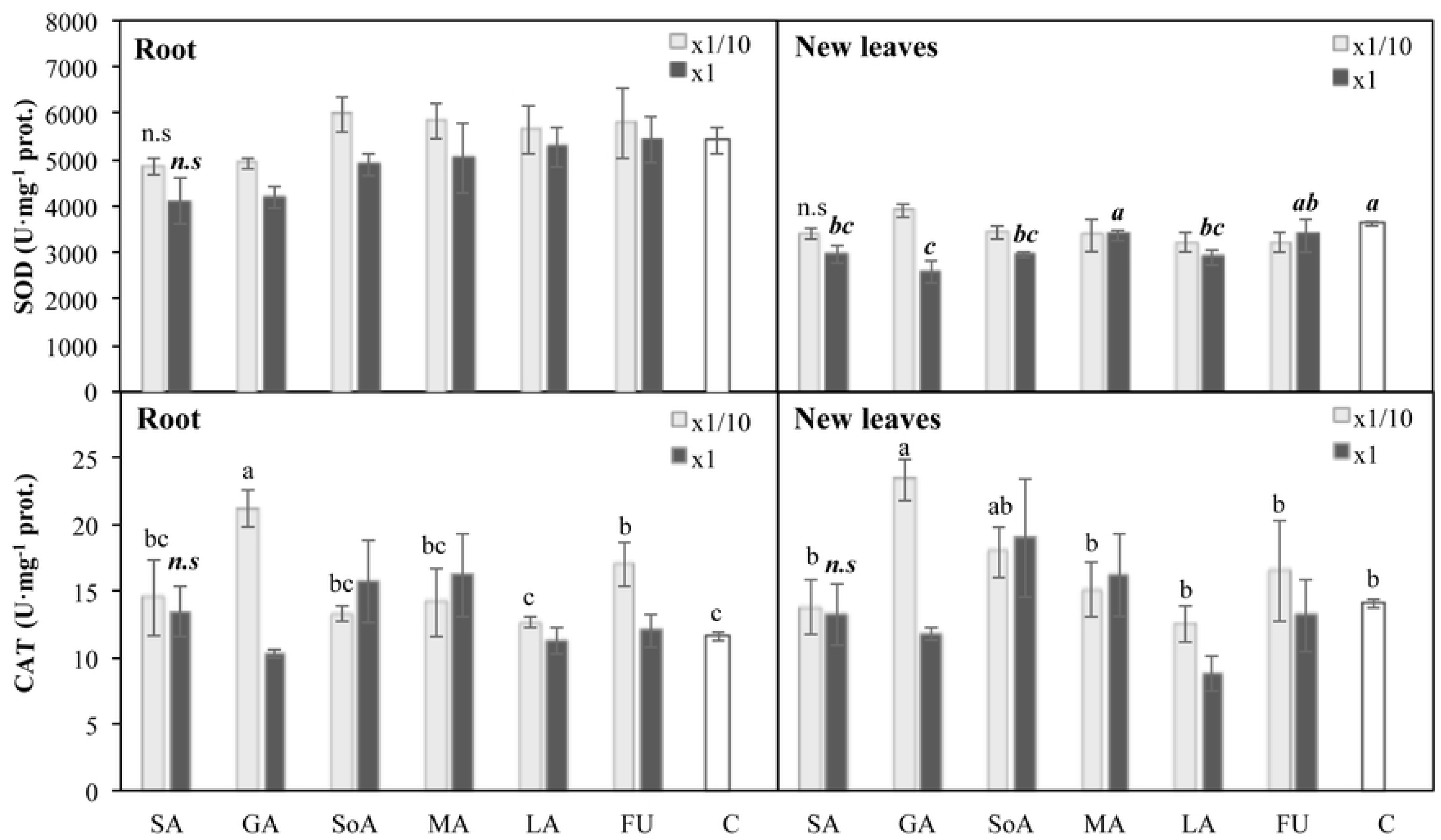
Oxidative stress indexes (SOD and CAT activity) of root and new leaves of tomato plants treated with organic compounds (salicylic acid (SA), gallic acid (GA), sodium alginate (SoA), mannitol (MA), laminarin (LA) and fucose (FU)) at two concentrations (x1/10 and x1) and after 6 days of Fe deficiency. A control (C) treatment without organic compound application was performed. Data are means ± SE (n=3). Significant differences between treatments (P < 0.10) are indicated by different letter (regular letters for x1/10; cursive bold letters for x1). Not significant differences between treatments are indicated by n.s.

## Discussion

Seaweed extract application may improve germination and plant growth development [36,37]. This promotion effect may be caused not only by phytohormones, but also by other compounds present in SWEs [5,38]. In fact, all organic compounds applied at different concentrations (x1/10, x1 and x10) in tomato seeds significantly increased the germination with respect to untreated control, except in the case of SA (x1 and x10) and SoA (x1). It has been reported that the positive effect of these compounds depends on the concentration applied. In an experiment with bean seedlings, the application of 0.1 mM SA (14 mg/L) inhibited germination and initial growth, but SA concentrations lower than 0,025 mM (3,4 mg/L) had a positive effect [39]. This study was in agreement with the obtained data, where the application of 47 and 470 mg/L SA (x1 and x10 respectively) inhibited the germination and the initial growth of tomato seedlings. Other example that showed the effect of concentration was the GA treatments, since the germination and the root length at day 3 increased as the concentration of GA applied decreased (470-47-4.7 mg/L). Similar results were found in a study that investigated the dose-response effect of GA on root growth in *Arabidopsis thaliana* at different concentrations 0, 150, 250, 500 and 1000 μM, where the highest concentrations of GA showed the lowest growth promotion (only 25%) [40]. SoA (x1/10 and x10) applications increased the germination, and also SoA (x1/10 and x1) promoted the root growth between 10 and 30% during 5 days in tomato seedlings. In concordance with the presented data, it has been reported that alginate derived oligosaccharides enhanced seed germination in maize [41], and increased the root growth in barley [42], lettuce [43] carrot and rice [44]. All MA treatments increased the germination and also MA (x1 and x10) enhanced, but not significantly, the root length until day 5 of growth. Contrary with our results, Johnson and Kane [45] indicated that MA application did not improve the germination in pine-pink seeds, and even very high concentration (7-9% (w/v)) inhibited the seed germination in celery plants by causing osmotic stress [46]. LA treatments enhanced the seed germination and also LA (x1/10 and x10) increased the root length, especially after 3 days of growth. Some authors indicated that LA might be used as a seed germination and plant growth accelerator in many plants [47]. FU treatments showed a very slight increase in seed germination, and except FU (x1/10), FU did not promote the root growth compared to the control. Stevenson and Harrington [48] applied FU in *Arabidopsis thaliana* seeds and showed a significant decrease in the hypocotyl and root length.

Regarding fresh weight, the lowest concentration (x1/10) increased the number of organic compounds that had a positive effect on biomass after six days of Fe deficiency. In fact, the concentration x1/10 of all organic compounds significantly increased the biomass of new leaves and roots (except SoA that only increased the root FW with the concentration x1). Moreover, GA, LA and FU at concentration x1/10 (4.7, 0.07 and 3.1 mg/L respectively) significantly increased the FW of all organs of the plant. Several studies reported positive effects on plant biomass under abiotic stress conditions after the application of SA, GA, SoA, MA, and LA, although none with respect to nutrient imbalance. Related to this, SA application at different concentrations, 0.5 mM (69 mg/L) in seeds of wheat [49], 100-300 mg/L sprayed in rosemary leaves [50] or 0.7-1.4 mM (96-193 mg/L) sprayed in sunflower [51] with high salinity showed a significant increase in FW. Applications in the nutrient solution of 1-2 mM (170-240 mg/L) GA in soybean under low temperatures [22] or applications of 0.75-1 mM (127-255 mg /L) GA in rice under salinity [20] increased the relative growth rate compared to the control. In the case of SoA, previous soaking of wheat seeds with 1000 mg/L alginate-derived oligosaccharide under cadmium toxicity [52] and 1000 mg/L alginate-derived oligosaccharide sprayed in rice plants under water stress [53], enhanced the FW. Application of 100 mM (1.82× 10^4^ mg/L) MA in the nutrient solution of wheat seedling with high salinity increased the root dry weight [12] and application of 15-30 mM (2.7×10^3^ – 5.4×10^3^ mg/L) MA by spray in maize leaves with high salinity increased the root and shoot dry weight [54]. Application of 25 mg/L LA in a growth medium under salt and heat stress showed an increase of *Arabidopsis thaliana* Col-0 FW [24]. As far as we know, the evaluation of FU application on plant biomass has been not described in the literature.

Iron deficiency induce root morphological alterations that results in a greater formation of root hairs [55] and secondary roots, a shorter length of the lateral roots [56], a decrease of the distance between secondary roots [57] and a thickening of root tips [56,58]. Under Fe deficiency, LA (x1/10) increased the length of secondary roots with respect to the untreated deficient control. Moreover, SA, GA, SoA, MA, FU (x1) and GA, LA, FU (x1/10) treatments increased the formation of root hairs under Fe deficiency, some of them (GA, MA (x1) and GA (x1/10)) even under sufficient conditions compared to the control. Related to this, some authors showed an increase of secondary roots formation after the application of seaweed extracts in *Arabidopsis thaliana* [5], grapevine [59] and strawberry [37]. This result suggests that the application of these organic compounds may contribute to the improvement of Fe availability, regulating the morphological adaptive responses of roots to Fe deficient conditions.

Despite several studies reported that the application of SWEs increased the chlorophyll content [5,26,60,61], others did not. For example, Carrasco-Gil et al. [62] did not obtained any change in the chlorophyll content after applying commercial SWEs to tomato plants after 7 days of Fe deficiency, and suggested that the application doses should be increased for attenuating chlorosis symptoms. In the present study, GA x1/10 (4.7 mg/L) and LA x1 (0.7 mg/L) significantly decreased the chlorophyll index of new leaves compared to the untreated control after three days of Fe deficiency. Contrary to these results, application of 60 mg/L GA in rice under healthy conditions [63] and 0.5 mg/L LA in tobacco under biotic stress [64] showed an increase of chlorophyll content or prevent chlorophyll depletion compared to the untreated control. However, FU x1/10 (3.1 mg/L) and FU x1 (31 mg/L) significantly increased the chlorophyll index of new leaves compared to the untreated control after five days of Fe deficiency, suggesting that fucoidans may increase the antioxidant activity reducing the chlorophyll degradation. As far as we know, no literature is available at this respect.

Iron deficiency enhanced the production of reactive oxygen species (ROS) in plants [65,66]. However, the role of ROS in Fe response regulation has not been well defined, and it may play multiple roles [67]. Plants have an enzymatic antioxidant system for scavenging the ROS excess and prevent damages to cells [68]. Superoxide dismutase (SOD), and catalase (CAT) are the first enzymes in the detoxification pathway and contain Fe in heme (CAT) and non-heme (Fe-SODs) form. Iron deficiency in plants increased total SOD activity (decreasing Fe-SOD and increasing CuZn-SOD and Mn-SOD), and reduced CAT activity since the synthesis of these enzyme is inhibited [66,69–71]. Several studies reported an increase of total SOD activity in leaves after the application of seaweed extracts in healthy plants [72], in Fe deficient plants [62] and in drought or water stressed plants [74,75]. In the present work, total SOD activity significantly decreased in Fe deficient tomato leaves after the application of SA, GA, SoA and LA (x1; 47, 47, 90 mg/L respectively) compared to untreated leaves. It could be possible that these compounds at concentration x1 balanced the ROS production (superoxide radical; O2^·-^), decreasing total SOD activity. Contrary to the results obtained, several authors reported an increase of total SOD activity after the application of GA and SoA in plants. Application of 1-2 mM (170-240 mg/L) GA in soybean grown in normal and cold stressed condition [22] and the application of 1000 mg/L alginate derived oligosaccharides in wheat grown in normal, drought and Cd stressed conditions [52,53], increased the SOD activity. In the case of SA, contradictory results have been found. Some studied showed an increase of total SOD activity in leaves after the application of 0.1-0.5 mM (14-69 mg/L) SA in root or leaves respectively of Fe deficient peanut [76] or 0.5 mM SA in soybean roots under arsenic toxicity [77]. However, other studies reported that 0.5 mM SA applied in maize plants under low temperatures did not affect the SOD activity [78,79], and even high concentrations of SA (2.5 mM; 345 mg/L) applied in wheat seedlings decreased total SOD activity [73]. On the other hand, CAT activity decreased in Fe deficient plants. However, application of SA, GA and FU x1/10 significantly increased the CAT activity in roots compared to Fe deficient control. Similar results were reported in salt stressed rosemary plants where the application of SA (100-300 mg/L) increased CAT activity [50]. The CAT enzyme catalyze the decomposition of hydrogen peroxide (H_2_O_2_) into oxygen and water [30]. Some authors reported a decrease of H_2_O_2_ in plants grown in salt [20] and cold [22] stressed condition after the application of GA suggesting an increase of CAT activity. Therefore, these phenolic compounds at concentration x1/10 may improve the antioxidant system enhancing the plant tolerance to Fe deficient stress. As far as we know, direct effects of fucoidan on plants have not yet been reported.

## Conclusions

In summary, the results of this research point out the importance of the concentration applied and the type of organic compounds present in a SWE in relation to its effectiveness to enhance the tolerance to iron deficiency (nutrient imbalance). The lowest concentration x1/10 of organic compounds showed the best results regarding growth promotion seedlings, fresh weight, secondary root elongation, chlorophyll content and CAT antioxidant activity. Moreover, from among all organic compounds evaluated, the phenolic compounds (salicylic acid and gallic acid), laminarin and fucose contributed in a greater extent to increase the tolerance to Fe deficiency in tomato plants. Although, it is necessary to carry out more studies in this regard, since it is possible that the effects of the algae extracts are not only due to the presence of discrete compounds, but the synergy produced by the interaction between them. It would also be of interest to test these compounds on other crops and substrates. In addition, experiments must be carried out to establish the optimal application times for these compounds, both in relation to the vegetative phase and the frequency of application. The achievement of these studies would be of great importance to establish greater control over the existing marine algae extracts, as well as to develop second generation algae extracts, designed with specific compositions for the needs of each crop.

## Contributions

Conceptualization: SCG, JJL. Metodology: SCG. Formal Analysis: SCG, RAM. Investigation: SCG, RAM. Writing-Original Draft: SCG, RAM. Formal analysis: SCG, RAM. Validation: LHA, JJL. Writing-Review and Editing: SCG, LHA, JJL. Funding Acquisition: LHA, JJL. Supervision: JJL.

## Abbreviations

SWE: seaweed extract
SA: salicylic acid
GA: gallic acid
SoA: sodium alginate
MA: mannitol
LA: laminarin
FU: fucose

